# Characterization of a Large Pericentric Inversion in Plateau Fence Lizards (*Sceloporus tristichus*): Evidence from Chromosome-scale Genomes

**DOI:** 10.1101/2020.03.18.997676

**Authors:** Ana M. Bedoya, Adam D. Leaché

## Abstract

Spiny lizards in the genus *Sceloporus* are a model system among squamate reptiles for studies of chromosomal evolution. While most pleurodont iguanians retain an ancestral karyotype formula of 2n=36 chromosomes, *Sceloporus* exhibits substantial karyotype variation ranging from 2n=22 to 2n=46 chromosomes. In this study, we present two annotated chromosome-scale genome assemblies for the Plateau Fence Lizard (*Sceloporus tristichus*) in order to facilitate research on the role of pericentric inversion polymorphisms on adaptation and speciation. Based on previous karyotype work using conventional staining, the *S. tristichus* genome is characterized as 2n=22 with 6 pairs of macrochromosomes and 5 pairs of microchromosomes with a large pericentric inversion polymorphism on chromosome seven that is geographically variable. We provide annotated, chromosome-scale genomes for two lizards located at opposite ends of a dynamic hybrid zone that are each fixed for different inversion polymorphisms. The assembled genomes are 1.84 to 1.87 Gb (1.72 Gb for scaffolds mapping to chromosomes) with a scaffold N50 of 267.5 Mb. Functional annotation of the genomes resulted in 65,417 annotated genes, 16,426 of which were deduced to have a function. We confirmed the presence of a 4.62 Mb pericentric inversion on chromosome seven, which contains 59 annotated coding genes with known functions. These new genomic resources provide opportunities to perform genomic scans and investigate the formation and spread of pericentric inversions in a naturally occurring hybrid zone.

## Introduction

Chromosomal rearrangements play important roles in adaptation, divergence, and speciation (Wellenreuther and Bernatchez, 2018). Pericentric chromosome inversions, which are classified as inversions that include a centromere, impose significant evolutionary and ecological constraints on genome evolution by trapping alleles on inverted chromosome segments. This greatly reduces recombination with non-inverted regions, and increases linkage disequilibrium within inverted regions (Kirkpatrick, 2010). Furthermore, when locally adapted alleles are located inside of an inversion, such as those that are ecologically relevant, the inversion can spread to fixation in a population (Kirkpatrick and Barton, 2006).

The phrynosomatid lizard genus *Sceloporus* is a diverse clade containing 108 species with a broad distribution across North America (Leaché et al., 2016). Differentiation in the fundamental number of chromosomes is hypothesized to be a factor responsible for driving the rapid diversification of the genus (Hall, 2009; Leaché and Sites Jr, 2009). While many species have the ancestral karyotype formula of 2n=36 chromosomes, *Sceloporus* exhibits substantial karyotype variation (ranging from 2n=22 to 2n=46), sex chromosome evolution, and large genome rearrangements (Leaché and Sites Jr, 2009). Existing genomic resources for *Sceloporus* include an annotated genome for *S. undulatus* (Westfall et al., prep), a *de novo* assembled shotgun genome for *S. occidentalis* and partial genomes (∼2.7% complete) for 34 other species (Genomic Resources Development Consortium et al., 2015).

Here, we present two annotated chromosome-scale genome assemblies for the Plateau Fence Lizard (*Sceloporus tristichus*), which is 2n=22 with 6 pairs of macrochromosomes and 5 pairs of microchromosomes, a karyotype formula shared with all member of the *undulatus* species group (10 species) (Leaché et al., 2016). An early study of the karyological differences within the group using conventional staining techniques identified a distinctive pericentric inversion polymorphism on chromosome seven that varies geographically (Cole, 1972). This inversion polymorphism produces distinctly recognizable variants classified by the position of the centromere (Cole, 1972), and individuals with heteromorphic pairs of distinct chromosome seven inversions can be found in a dynamic hybrid zone (Leaché and Cole, 2007). The hybrid zone occurs in Arizona’s Colorado plateau at the ecotone between Great Basin Conifer Woodland and Grassland habitats (Leaché and Cole, 2007). Temporal comparisons of clinal variation in kayotypes, morphology, mitochondrial DNA, and SNPs suggests that the hybrid zone is moving, possibly as a response to habitat modification (Leaché and Cole, 2007; Leaché et al., 2017). The new genomic resources reported here provide a framework for increasing our understanding of the formation and spread of large chromosome inversions across broad temporal and spatial scales.

## Materials and Methods

### Sampling, Library Preparation, and Sequencing

We sequenced two female specimens of *Sceloporus tristichus* collected from a hybrid zone in Navajo County, Arizona, USA in 2002. The two individuals were karyotyped to determine the position of the centromere on chromosome seven in an earlier study (Leaché and Cole, 2007). One specimen (NCBI BioSample ID SAMNxxxxxxx; voucher specimen AMNH 153954) is from the northern end of the hybrid zone (Holbrook; HOL), and is submetacentric at chromosome seven (a nonmedial centromere that is closer to the middle than to either end of the chromosome). The second specimen (NCBI BioSample ID SAMNxxxxxxx; voucher specimen AMNH 154032) is from the southern end of the hybrid zone (Snowflake; SNOW), and is telocentric at chromosome seven (a terminal centromere that is close to the end of the chromosome).

Genome sequencing was performed by Dovetail Genomics (Santa Barbara, California, USA). High molecular weight genomic DNA was extracted from flash frozen liver tissues and validated for large fragment sizes (50kb-100kb) using pulsed field gel electrophoresis. 10X Chromium libraries were prepared using the Chromium Genome Reagent Kit (10x Genomics, Pleasanton, CA) followed by Illumina HiSeqX sequencing (150bp paired-end reads). Chicago (Putnam et al., 2016) and Hi-C libraries (Belton et al., 2012) were prepared at three gigabase increments of the total genome size. Quality control for these libraries was performed by mapping reads (75bp paired-end MiSeq reads) to draft 10x Supernova assemblies. Finally, both libraries were sequenced on an Illumina HiSeqX (150bp paired-end reads). Raw sequencing reads are deposited in the NCBI Sequence Read Archive (SRAxxxxx; SRAxxxx).

### Genome Assembly and Annotation

For each individual, a draft genome was *de novo* assembled from 709.35 million 10X Chromium sequence reads using the *SuperNova* assembly pipeline. Chromosome-scale scaffolding was achieved by mapping the Chicago and Dovetail Hi-C libraries back to the 10x Supernova assembly (Weisenfeld et al., 2017) using the *HiRise* software pipeline (Putnam et al., 2016).

Functional annotation of the two genomes was conducted with Funnanotate v1.7.2. (Palmer and Stajich, 2019). Briefly, Funannotate aligns raw RNASeq datasets to a genome sequence with minimap2 and assembles them using Trinity (Grabherr et al., 2011). Such assemblies are used along with PASA predictions (Haas et al., 2003) to build consensus gene model predictions, and train Augustus (Stanke et al., 2004) and GeneMark-ES/ET (Lomsadze et al., 2005). We used the Tetrapoda BUSCO (Benchmarking Universal Single-Copy Orthologs) dataset for gene prediction (Simão et al., 2015). Finally, Funnanotate performs genome functional annotations using the outputs of PFAM (Finn et al., 2016), MEROPS (Rawlings et al., 2018), InterProScan5 (Jones et al., 2014), eggNOG-mapper (Huerta-Cepas et al., 2017), and Phobius (Käll et al., 2004). We run the pipeline for each of our two *HiRise* genome assemblies and included a raw RNASeq dataset prepared from skeletal muscle of an adult female of *S. undulatus* (BioSample SAMN06312743) to build consensus gene model predictions. Annotation was performed on the 24 longest scaffolds, which included 92.3% and 93.3% of the total length of the *HiRise* assemblies (Table 1). All assemblies and functional annotation files are available at Dryad (doi/drayd.org/xxxxx).

**Table 1:** Summary statistics for the *HiRise* genome assemblies. Sequence lengths are given in megabases.

| Metric | HOL | SNOW |
| --- | --- | --- |
| Assembly Size | 1,870.61 | 1,844.30 |
| Scaffolds >10Kb | 25,615.68X | 32,060.90X |
| L50 <sup>a</sup> | 3 | 3 |
| N50 <sup>b</sup> | 269.061 | 267.474 |
| L90 <sup>c</sup> | 15 | 10 |
| N90 <sup>d</sup> | 17.367 | 36.595 |
| Chromosome 1 | 366.851 | 366.429 |
| Chromosome 2 | 304.053 | 319.091 |
| Chromosome 3 | 269.060 | 267.474 |
| Chromosome 4 | 249.456 | 247.712 |
| Chromosome 5 | 173.679 | 170.938 |
| Chromosome 6 | 115.627 | 114.258 |
| Chromosome 7 | 38.601 | 68.322 |
| Chromosome 8 | 27.143 | 48.193 |
| Chromosome 9 | 26.412 | 38.926 |
| Chromosome 10 | 23.766 | 36.595 |
| Chromosome 11 | 22.414 | 14.995 |
| Top 24 scaffolds | 1,727.963 | 1,721.029 |

<sup>a</sup>L50 = Minimum number of scaffolds that make up 50% of the total assembly length.
<sup>b</sup>N50 = Minimum scaffold size of fragments that make up 50% of the total assembly length.
<sup>c</sup>L90 = Minimum number of scaffolds that make up 90% of the total assembly length.
<sup>d</sup>N90 = Minimum scaffold size of fragments that make up 90% of the total assembly length.

### Pericentric Inversion Polymorphism

To identify the locations of inversions on chromsome seven between SNOW and HOL we used progressive Mauve v.2.4.0 (Darling et al., 2004). The program was run with default “seed families” and default values for all other parameters. We determined the identity of large scaffolds by comparing *S. tristichus* assemblies to the assembled chromosome-scale genome of the closely related species *S. undulatus* (Westfall et al., prep).

## Results and Discussion

### Assembly and Annotation

10X SuperNova assemblies included 31,453 and 30,454 scaffolds at 53.06X and 53.88X coverage, with a total assembly length of 1870.17 Mb and 1843.78 Mb. In turn, the final *HiRise* assemblies resulted in 27,095 and and 25,281 scaffolds at 25,615X and 32,060X coverage, adding to a total length of 1,870.61 Mb and 1,844.30 Mb for HOL and SNOW, respectively. Genome assembly statistics are provided in Table 1 and in the Supplementary Material. The top 11 scaffolds account for 86.44% and 91.79% of the total length of the HOL and SNOW assemblies, and comparisons to the *S. undulatus* genome confirmed that these scaffolds are the 11 chromosomes. The top six scaffolds are on average 206 Mb longer than the subsequent five, which is expected given that *S. tristichus* has 6 macrochromsomes.

Functional annotation resulted in 65,417 annotated genes, 16,426 of which were deduced to have a function by Funannotate (Dryad DOI/dryad/xxxx.xxxx). The remaining genes were annotated as hypothetical proteins. Chromosome seven spanned 2,730 annotated genes for HOL, whereas 3,353 were identified for SNOW (Dryad DOI/dryad/xxxx.xxxx). The higher number of annotated genes in SNOW is the result of a higher quality and more complete assembly compared to HOL as interpreted from the assembly summary statistics in Table 1.

### Pericentric Inversion Polymorphism

Alignment of chromosome seven with Mauve resulted in the detection of a 4.62Mb pericentric inversion (Figure 1). This chromosomal rearrangement distinguishes populations of *S. tristichus* located at opposite ends of a dynamic hybrid zone and is expected to reduce recombination in hybrids with heteromorphic paris of chromosome seven, which are found at the center of the hybrid zone (Leaché and Cole, 2007).

**Figure 1:**
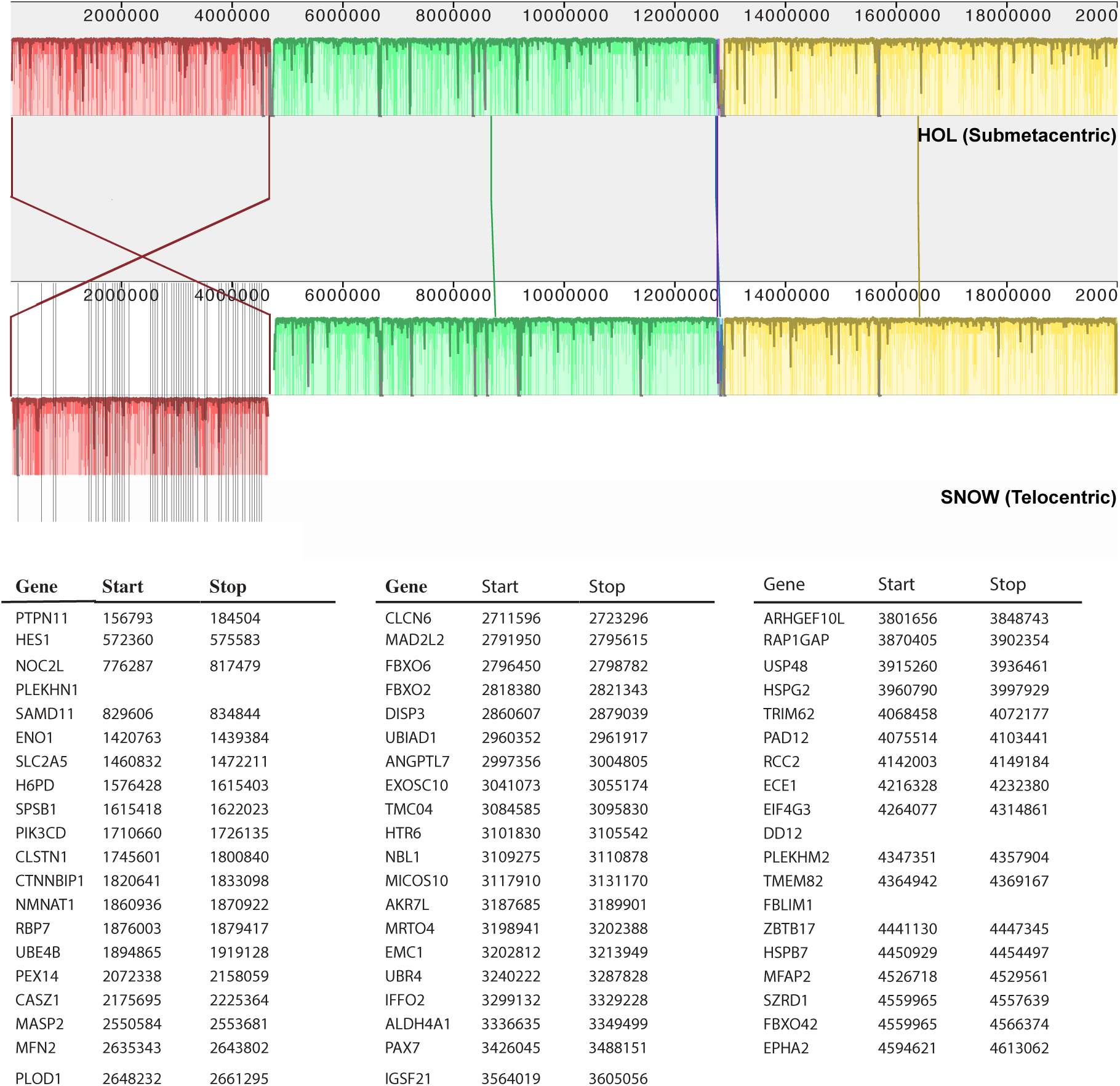
Pericentric inversion polymorphism on chromosome seven for the two *Sceloporus tristichus* lizard genomes and position of the coding genes across the inversion. Vertical bars indicate the position of the coding genes. Only the first 20 Mb are shown.

A total of 59 annotated genes with known function are found within the region that spans the inversion (Figue 1 and Supplemental Materials). Whether any of these genes are involved in local adaptation or fixation of the inversion remains to be examined. The large size of this inversion makes it more likely that it would span multiple loci involved in local adaptation. However, the large size also makes it more likely to suffer meiotic costs that could reduce hybrid fitness (Kirkpatrick and Barton, 2006).

These new genomic resources provide a starting point for understanding how a pericentric inversion polymorphism could cycle in frequency in a dynamic hybrid zone that is moving through time. Identifying the location, size, and gene content of this inversion polymorphism provides the necessary tools for investigating the role of inversions on local adaptation and speciation.

## Supplementary Material

Supplementary materials are available at Genome Biology and Evolution online.

## Acknowledgments

We are thankful to the University of Washington for financial support through the Bridge Funding program awarded to ADL. We also thank Tonia Schwartz and Damien Waits at Auburn University for useful feedback on Funannotate.

## Author Contributions

ADL conceived the study, AMB analyzed the data, and both authors wrote the manuscript.

