## Supplementary Materials for "Characterization of a Large Pericentric Inversion in Plateau Fence Lizards (*Sceloporus tristichus*): Evidence from Chromosome-scale Genomes"

This PDF file includes: Tables S1 to S2

Table S1: Summary statistics for the 10X Chromium, Chicago, and Dovetail Hi-C data generated in this study.

| Individual | Metric | 10X Chromium | Chicago | Dovetail Hi-C |
|---|---|---|---|---|
| HOL | No. Scaffolds (>10Kb) | 31,453 | 27,316 | 27,095 |
| | Assembly size | 1870.17 | 1870.59 | 1870.61 Mb |
| | Coverage (Scaffolds >10Kb) | 53.06 | 211.07X | 25615.68X |
| | L50 | 19 | 26 | 3 |
| | N50 | 28.61 | 22.866 Mb | 269.061 Mb |
| | L90 | 120 | 133 | 15 |
| | N90 | 1.129 Mb | 1.508 Mb | 17.367 Mb |
| SNOW | No. Scaffolds (>10Kb) | 30,453 | 25,459 | 25,281 |
| | Assembly size | 1843.78 Mb | 1844.28 Mb | 1844.3 Mb |
| | Coverage (Scaffolds >10Kb) | 53.88 | 198.38X | 32060.90X |
| | L50 | 14 | 15 | 3 |
| | N50 | 29.051 Mb | 30.900 Mb | 267.474 Mb |
| | L90 | 129 | 95 | 10 |
| | N90 | 1.199 Mb | 2.267 Mb | 36.595 Mb |

Table S2: Fully annotated coding genes located in the pericentric inversion in chromosome seven in *Sceloporus tristichus* from Holbrook(HOL) and Snowflake(SNOW).

| Gene | Function |
|---|---|
| PTPN11_1 | protein tyrosine phosphatase* |
| HES1 | oxysterol-binding protein hes1 |
| NOC2L | Nucleolar complex protein 2* |
| PLEKHN1 | Pleckstrin y domain-containing N member 1* |
| SAMD11 | Sterile alpha motif domain-containing protein 11 |
| ENO1 | phosphopyruvate hydratase |
| SLC2A5 | Solute carrier 2 |
| H6PD | 6-phosphogluconolactonase |
| SPSB1 | SPRY domain-containing SOCS box protein 1 |
| PIK3CD | Phosphatidylinositol 4 |
| CLSTN1 | Calsyntenin-1 |
| CTNNBIP1 | Beta-catenin-interacting protein 1* |
| NMNAT1 | Nicotinamide/nicotinic acid mononucleotide adenylyltransferase 1 |
| RBP7 | Retinoid-binding protein 7 |
| UBE4B | Ubiquitin conjugation factor E4 B |
| PEX14 | Kinesin-like protein kif1b |
| CASZ1 | Zinc finger protein castor 1* |
| MASP2 | Mannan-binding lectin serine protease 2 |
| MFN2 | Mitofusin-2 |
| PLOD1 | Procollagen-lysine |
| CLCN6 | Chloride transport protein 6 |
| MAD2L2 | MAD2 mitotic arrest deficient-like 2 |
| FBXO6 | F-box only protein 6 |
| FBXO2 | F-box only protein 2 |
| DISP3 | Protein dispatched 3 |
| UBIAD1 | UbiA prenyltransferase domain-containing protein 1 |
| ANGPTL7 | Angiopoietin- protein 7 |
| EXOSC10 | Exosome component 10 |
| TMCO4 | Transmembrane and coiled-coil domain-containing protein 4 |
| HTR6 | 5-hydroxytryptamine receptor 6 |
| NBL1 | Borealin |
| MICOS10 | MICOS complex subunit* |
| AKR7L | Aflatoxin B1 aldehyde reductase member 2 |
| MRTO4 | mRNA turnover 4 |
| EMC1 | DUF1620 super |
| UBR4 | E3 ubiquitin-protein ligase ubr4 |
| IFFO2 | Intermediate filament orphan 2 |
| ALDH4A1 | Delta-1-pyrroline-5-carboxylate dehydrogenase |
| PAX7 | Paired box protein Pax-7 |
| IGSF21_1 | Immunoglobulin super member 21 |
| ARHGEF10L | guanine nucleotide exchange factor 10-like protein |
| RAP1GAP | Rap1 GTPase-activating protein 1 |
| USP48 | Ubiquitin carboxyl-terminal hydrolase 48 |
| HSPG2 | Basement membrane-specific heparan sulfate proteoglycan core protein |
| TRIM62 | E3 ubiquitin-protein ligase trim62 |
| PADI2 | Protein-arginine deiminase type-2 |
| RCC2 | Protein rcc2* |
| ECE1 | Endothelin-converting enzyme 1 |
| EIF4G3 | Eukaryotic translation initiation factor 4 gamma |
| DDI2 | cyanamide hydratase* |
| PLEKHM2 | Pleckstrin y domain-containing M member 2 |
| TMEM82 | Transmembrane protein 82 |
| FBLIM1 | Filamin-binding LIM protein 1* |
| ZBTB17 | Zinc finger and BTB domain-containing protein 17 |
| HSPB7 | Heat shock protein beta-7 |
| MFAP2 | Microfibrillar-associated protein 2* |
| SZRD1 | SUZ domain-containing protein 1* |
| FBXO42 | F-box only protein 42 |
| EPHA2 | Ephrin type-A receptor 2 |

(*) Coding genes only annotated in one assembly.
